# Teachers’ choice of content and consideration of controversial and sensitive issues in teaching of secondary school genetics

**DOI:** 10.1101/350710

**Authors:** Tuomas Aivelo, Anna Uitto

## Abstract

Science education strives to increase interest in science and facilitate active citizenship. Thus, the aspects of personal and societal relevance are increasingly emphasised in science curricula. Still, little is known about how teachers choose content for their teaching, although their choices translate curricula to teaching practice. We explored how teachers choose genetics content and contexts for biology courses on cells, heredity and biotechnology by interviewing ten Finnish upper-secondary school teachers. We specifically studied how the teachers described teaching on genetically modified organisms, hereditary disorders, and complex human traits as teachers have different amounts of freedom afforded by curricula in choosing contents and contexts on these themes. We analysed interviews with theory-guiding content analysis and found consistent patterns in teachers’ perceptions of the main themes in genetics teaching, teacher inclinations towards teaching genetics in human context and perceptions of students’ interest in different topics. These patterns, which we call emphasis of content in genetics teaching could be classified to *Developmental*, *Structural* and *Hereditary.* Teachers with *Developmental* emphasis embraced teaching genetics in a human context, while teachers with a *Structural* emphasis avoided them. In general, teachers justified their choices by national, local school, and personal factors. While teachers mentioned that societal and personal contexts are important, at the same time teachers never framed the main themes in genetics with these contexts. We conclude that how teachers handle issues of societal or personal relevance should be emphasized.

## Introduction

Curriculum articulates learning goals in school and guides teaching. In Finland, the national curricular goals are managed by legislation: the municipal authorities have an autonomy to provide and organise education at the local level and teachers are valued as experts who are able to develop and implement the school-specific curriculum (Niemi, Toom, & Kallioniemi, 2012). Thus, the curriculum allows teachers a remarkable responsibility and freedom to implement education in emphasizing the contents of upper-secondary school courses (Finnish National Board of Education, FNBE, 2003; Niemi et al., 2012). While curricular development and its effectiveness have been studied extensively (e.g., Hargreaves, Lieberman, Fullan, & Hopkins, 2010; Niemi et al., 2012), there has been less research produced on how teachers choose course content for their teaching.

To substantial extent, teacher beliefs guide how teachers value different aspects of knowledge and how much they emphasise different content (Cheung & Wong, 2010; Cronin-Jones, 1991; Haney, Czerniak, & Lumpe, 1996). Teachers’ ‘personal knowledge’ is not static, but is formed through everyday experiences and formal schooling, including teacher education; it is further moudled in continuing professional education (Gess-Newsome & Lederman, 1995; Henze, Van Driel, & Verloop, 2007; Van Driel, Beijaard, & Verloop, 2001). Thus, this personal knowledge shapes teaching to a large extent (Hashweh, 1987, 2005; van Driel, Bulte, & Verloop, 2008). Nevertheless, personal knowledge often manifests through rules-of-thumb, rather than formal design (Wieringa, Janssen, & van Driel, 2011). Consequently, curricular change can cause truly little change in teaching approaches if underlying beliefs about content and the best suited methods to teach content do not change in teachers’ practice (Cohen & Yarden, 2009; Tidemand & Nielsen, 2017).

The main tools for content selection are the available teaching materials, especially textbooks (Remillard & Bryans, 2004; Shawer, 2017; Spiegel & Wright, 1984). Teachers have a conflict between following the textbooks’ content and using them critically, as teachers seem to understand critical reading of texts as distancing themselves from the text (Loewenberg Ball, Feiman-Nemser, Ball, & Feiman-Nemser, 1988).

### Controversial and sensitive issues in teaching

According to the definition used by Oulton, Dillon, and Grace (2004), we define controversial issues as issues on which groups within society hold differing views based on different sets of information or different interpretations from the available information due to their worldview, such as different value systems. Sometimes controversial issues may be resolved by acquiring additional information, but not always. For sensitive issues, we follow the ideas of Rowling (1996), who suggests that the distinction between sensitive and controversial issues seem to be that sensitive issues are connected to emotionality and the involvement of the individual. Sensitivity can arise from political, religious, cultural, personal or gender sources, but in comparison to controversial issues, which by definition usually work on a societal level, sensitive issues are more personal.

Real world applications of science commonly involve controversial issues; however, teachers are generally poorly prepared for teaching controversial issues (Oulton, Dillon, & Grace, 2004; Oulton, Day, Dillon, & Grace, 2004). Furthermore, less experienced teachers do not seem to select topics that could be upsetting to students (Hess, 2008; Phillips, 1997). Controversial and sensitive issues may appear when teaching different contents of biology, especially in the framework of socio-scientific issues (Lederman, Antink, & Bartos, 2014; Lewis & Leach, 2006; Zeidler, Walker, Ackett, & Simmons, 2002). Nevertheless, it is known that some teachers are unwilling to use this framework in their teaching (Lazarowitz & Bloch, 2005; Lee, Abd-El-Khalick, & Choi, 2006). Several reasons have been proposed to explain this phenomenon, including limitations of the curriculum or assessment techniques, teachers’ pedagogical competence and their lack of support for the merits of SSI discussions as pertinent to specific subjects (Bryce & Gray, 2004; Gray & Bryce, 2006; Lewis & Leach, 2006; Newton, Driver, & Osborne, 1999).

Arguably teacher beliefs are important factors that affect whether teachers embark on discussions of controversial issues (Cotton, 2006). Most of the research on teaching controversial issues has largely been in history and social science classes (i.e., Hess, 2008; Oulton et al., 2004a; Oulton et al., 2004b) while sensitive issues are discussed in health and physical education (Lynagh, Gilligan, & Handley, 2010; Rowling, 1996). Science subjects have been less studied, even though they do not lack controversial or sensitive topics (Leonard, 2010; Levin & Lindbeck, 1979; Owens, Sadler, & Zeidler, 2017). In discussing controversial issues, teachers have mentioned problems in beginning and maintaining discussions, dealing with students’ “lack of knowledge”, insufficient teaching time, and scarcity of resources (Dawson & Venville, 2008; Dawson & Taylor, 2000; Hand & Levinson, 2012; Kuş, 2015; Reiss, 1999). One of the solutions should be finding ways to increase teachers’ confidence to implement teaching societally relevant issues, even though they could be controversial (Hofstein, Eilks, & Bybee, 2011).

Teachers’ beliefs seem to relate to their teaching in several ways. For example, their own understanding whether they are experts in biology versus experts in discussing human genetics seems to be one of the central problems (Pedretti, Bencze, Hewitt, Romkey, & Jivraj, 2008; Tidemand & Nielsen, 2017). Teachers’ content and context choice also have been found to affect also students’ learning: for example, in a Swedish study, upper-secondary school students majoring in science used few justifications from ethics or morality when discussing GMOs (Christenson, Chang Rundgren, & Zeidler, 2014).

#### Genetics content and curricular development

Genetics in the secondary school biology curriculum has been emphasised in recent years, as the progress in both basic science of genetics and its technological applications has been rapid. This is likely to lead to both curricular renewal and the requirement for continued teacher development. Choosing content for genetics courses is challenging for teachers and that difficult process illustrates a teacher’s perspective on teaching genetics.

There have been a few endeavours to outline the core (conceptual) contents of genetics on different levels of lower and upper secondary school curricula. Stewart, Cartier and Passmore (2005) outlined that a basic understanding of genetics requires understanding three basic models: genetic (i.e., Mendelian inheritance patterns), meiotic (i.e., chromosome segregation and assortment) and biomolecular (i.e., the genotype-to-phenotype process). This was in turn refined by Duncan et al. (2009) who added environment as a context and outlined their learning progression around two big ideas: 1) “All organisms have genetic information that is universal and specifies the molecules that carry out the functions of life. While all cells have the same information, cells can regulate which information is used (expressed).” and 2) “There are patterns of gene transfer across generations. Cellular and molecular mechanisms drive these patterns and result in genetic variation. The environment interacts with our genetic makeup leading to variation.” Furthermore, in their Delphi study of genetic literacy, Boerwinkel et al. (2017) added a difference between somatic and germ line and polygenic inheritance to previous core contents and also emphasised also sociocultural and epistemic knowledge. In general, the diversification of core contents in genetics education from Mendelian genetics to polygenic traits seems to mirror the change in gene research emphasis from quantitative genetics to genomics and whole-genome sequencing. Combining the insights of all this research could help teachers gain more confidence in choosing content for genetics courses.

#### Genetics in the Finnish upper-secondary school biology curriculum

In Finland, approximately half of each age class enter general upper secondary schools, which aims to both provide general knowledge required for an active participation in society and prepare students for further education on the tertiary level and working life. Finnish biology teachers at upper secondary level have at least a Masters level degree, including one-year of studies in teacher education and one year of biology study (see Niemi et al., 2012).

Finnish curricula tend to leave substantial freedom for teachers to interpret the educational aims and develop multiple methods to implement curricula. Finnish teachers plan teaching according to local curricula, which are formulated by the education providers and schools based on the national core curriculum for general upper-secondary schools (FNBE, 2003) The core content pertaining to genetics is primarily limited to two courses: Cells and heredity (BI2), which is mandatory for all students and Biotechnology (BI5), which is an optional course in the biology curriculum (Table 1).

**Table 1:**
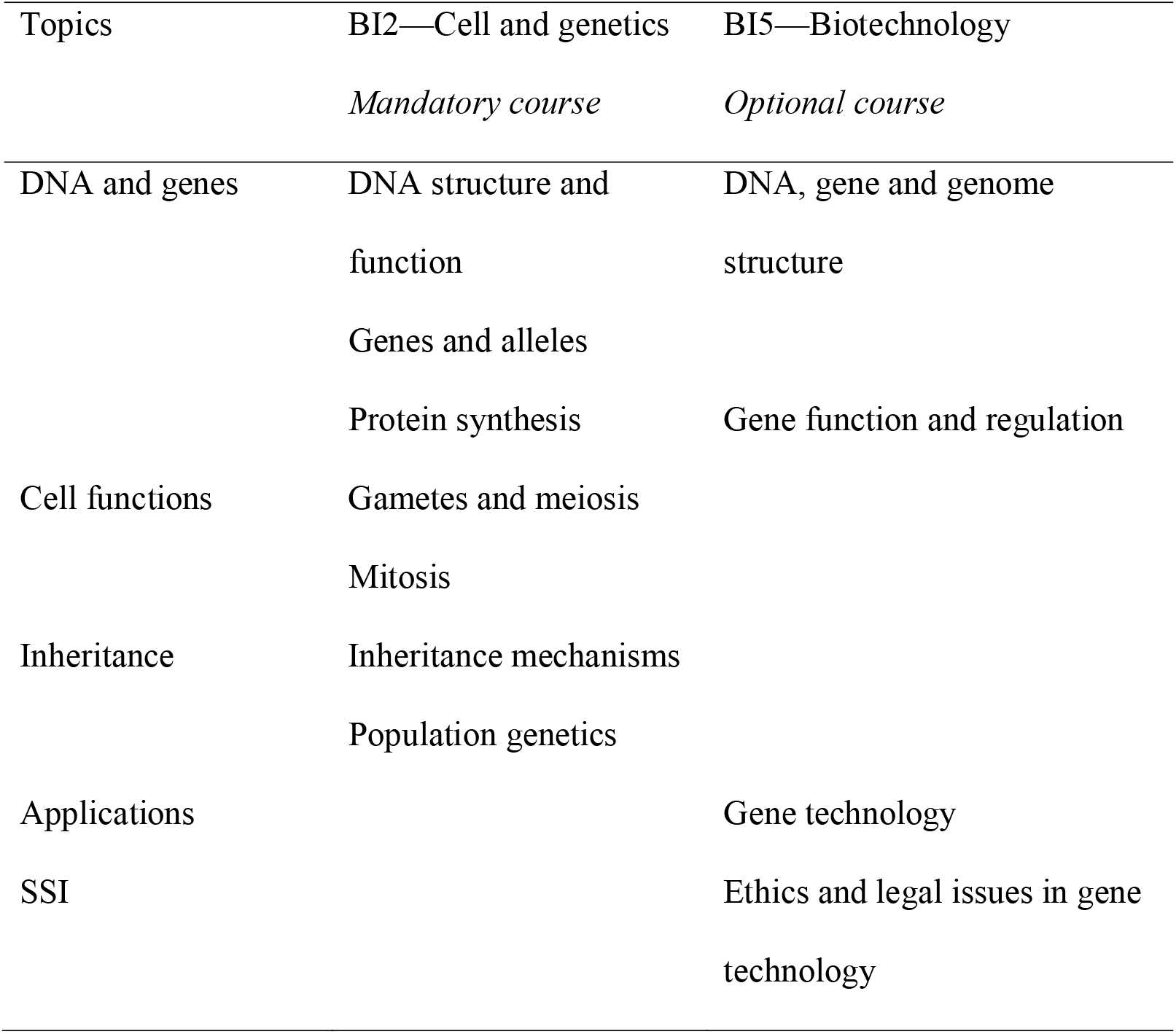
The Core content described in the Finnish national curriculum with selected parts from courses BI2 and BI5

While teachers’ practices and attitudes towards different teaching approaches and methods have been widely studied in science education (e.g., Lederman & Abell, 2014), there is far less research on what contents and examples teachers choose for their teaching and how they justify their choices in an upper-secondary school biology course. Because many contents of secondary school genetics can be taught in a human context, such as inheritance of genetic disorders, we were also interested in how teachers perceive controversial and sensitive issues during the biology lessons. As the Finnish upper secondary school curriculum provides ample freedom for teachers to adopt the most suitable teaching methods and biology teachers are generally educated broadly in different fields of biology, we were able to explore the curricular genetics contents the teachers emphasise in their teaching and what are teachers’ perceptions of controversial and sensitive issues are their teaching.

Our research questions were:

1. What do teachers suggest as the main contents of genetics teaching in the upper secondary school in biology?
2. How teachers argue for their use of human-related contexts in genetics teaching, specifically in themes of GMOs, human hereditary disorders, and complex human traits?
3. What kind of controversial or sensitive issues do teachers consider when teaching upper secondary school genetics in general, and specifically, the subjects of GMOs, human hereditary disorders, and complex human traits?

## Methods

### Overall study design

Our research design was a qualitative case study. We conducted open-ended semi-structured interviews with 10 upper-secondary high school biology teachers from various schools from Southern and Western Finland between 2015 and 2016 (see Table 2). Teachers were selected purposively to reflect a variation in experience, gender, type of school and geographical location to access different teachers with knowledge about upper secondary school biology education. All teachers studied biology as a major subject in their university master’s degree. Additionally, we collected diary data and other teaching materials from teachers about how they actually teach genetics.

**Table 2:** Summary of interviews of ten interviewed teachers. For details, see the supplemental material.

| Teacher | Gender | Central theme of teaching | Examples of Human Mendelian traits | Examples of Complex human traits | Perceived controversial or sensitive issues | Perceived student interest | Genetics content emphasis | Teacher concurrence with analysis |
| --- | --- | --- | --- | --- | --- | --- | --- | --- |
| A | Female | Development of phenotype | Disorders, lactose intolerance | Life style diseases | None | Artistic talent, epigenetics | Developmental | Yes |
| B | Male | Development of phenotype | Disorders, blood groups, tongue roll | Height, skin colour, talent | None | Epigenetics, talents, monohybrid crosses | Developmental | Yes |
| C | Female | Inheritance and continuity | Disorders | Eye color, **generally avoid** | Intelligence | Gene tests | Hereditary | Yes |
| D | Male | Inheritance and continuity | Disorders | Height, skin colour | Human race-related | Medical genetics | Hereditary | No answer |
| E | Male | The structure and function of genes | Eye colors, **generally avoid** | Shoe size, **generally avoid** | Intelligence | Mono- and dihybrid crosses | Structural | Yes |
| F | Female | Development of phenotype | Disorders, tongue roll, widow's peak | Height | None | Challenging contents | Developmental | No; Structural |
| G | Female | The structure and function of genes | Tongue roll, eye colors, **generally avoid** | Height, hair coloration, **generally avoid** | Intelligence, talent, genetic disorders | Mono- and dihybrid crosses | Structural | Yes |
| H | Female | Inheritance and continuity | Tongue roll, ear lobe | Skin color, generally avoid | None mentioned | Inheritance patterns | Hereditary | Yes |
| I | Male | Development of phenotype | Disorders | Stress reaction, intelligence | None | Sex-related traits | Developmental | No; Structural |
| J | Female | Development of phenotype | Disorders, eye colour, ear lobe | Height, skin colour | Developmental disorders | Musicality, own complex traits | Developmental | No; Hereditary |

We asked teachers’ for their perceptions on their teaching in the framework of Finnish upper secondary school biology and asked them to think about both compulsory BI2 and optional BI5 courses. The main contents of genetics teaching were asked in this context (Research Question 1). Furthermore, we asked teachers specifically how they teach and what examples they use on three different human-related contexts: genetically modified organisms (GMOs), human hereditary disorders and complex human traits, such as intelligence. These contexts function as a gradient on how much freedom teachers have to choose what contents and contexts they teach. The first context, GMOs, is explicitly mentioned in the national core curriculum and teachers must discuss the ethics of GMOs. The second context, hereditary disorders, is not mentioned in the curriculum, but most known examples of Mendelian genetic traits in a human context are hereditary disorders. Thus, teachers can avoid this context, but it is difficult. Thirdly, such complex human traits as intelligence are not mentioned in the curriculum but this context can be used to discuss polygenic inheritance. Thus, teachers can easily avoid this context and not use it all in their teaching. We asked teachers to explain their reasons why they use these contexts or why they would not use them (RQ2). The idea behind using this gradient of teacher freedom in relation to the curricula is to gauge how teachers relate their view on what is important to learn in genetics to their own choices in contexts and contents. Lastly, we asked teachers whether they perceived any controversial or sensitive issues in their genetics teaching in general or specifically in these three contexts (RQ3).

### Interviews

The interviews lasted from 40 minutes to 1 hour and 32 minutes. Teachers were asked: a) what they perceive as the most important contents and contexts in genetics, b) how they acquire knowledge for teaching and c) what examples they use during the two courses, the BI2, for Cells and heredity course for all students, and BI5, an optional Biotechnology course (Table 1). We specifically asked how teachers teach the topic of GMOs in the BI5 course and what kind of examples of human genetics they use in courses BI2 and BI5.

Our aim was to learn how teachers justify their content and context choices in genetics teaching. We used a theory-guided content analysis to categorize the data in a six-stage process by following the ideas of abductive analysis laid out by Timmermans and Tavory (2012): 1) we transcribed the interviews; 2) we coded the transcripts one sample at a time by looking for our units of analysis: a) which subject matter teachers thought was the most crucial and which could be left aside, b) how they argued for including or excluding certain course content and c) how they described what they believe students feel important; 3) beginning from the first sample, we named concepts arising from the grouped codes and after each sample, recursively performed stages 2 and 3 for previously-coded samples (which would correspond to initial analysis as per Charmaz (2003); 4) after initial samples were coded and concepts named, we integrated the categories through a focused analysis; 5) we contrasted the teachers to each other to understand the connections among categories, and 6) we refined the model. We used the R (R Core Team, 2013) package RQDA (Huang, 2017) for the analysis.

We contrasted the emerging codes with the assumption that teachers’ content and context choices are guided by the national and local curriculum, teaching materials and teachers’ personal knowledge. When coding content choices, three distinct groups emerged: monohybrid crosses in humans, polygenic properties of humans and GMOs. Within these three groups, we coded on later recursions all the mentions of the issues the teachers described that a) they use in teaching, b) they avoid using in teaching, c) the topics in which the students express interest and d) topics in which the students express no interest in. We then simplified authentic expressions in the open codes to a combination that would describe general-level biological phenomena, such as evolution, inheritance or gene expression. After half of the samples were coded, selective coding was used to delimit the coding process. Purposive sampling fitted well this research approach as our data becamerather rapidly saturated: by the ninth sample, there was no new information useful for the category formation. After the analysis, we asked teachers whether they agreed with our analysis of the emphasis of their teaching.

### Trustworthiness

To assess the connection between descriptions that the teachers gave of their teaching during the interviews and their actual teaching, we asked the teachers to keep a diary of their teaching after the interviews. We suggested that for each lesson the teachers record the topics, teaching methods, textbook chapters and exercises that were discussed and about which topics the students asked questions or needed clarification. Additionally, teachers who had ready-made lecture slides sent those to us. An outside observer and first author classified diaries and other materials based on the previously formed classifications (Table 2), which allowed us to compare teachers’ interviews and their actual teaching.

We continuously evaluated the trustworthiness of our study in several ways (Morrow, 2005). During the category formation, we looked for disconfirming data and assessed data saturation. The credibility was also enhanced by continuous discussion and revision of the meanings and data coding during the categorization by the first author and the transcriber, who was a sociolinguist. Transferability was improved by a rich description of the research process in the form of an audit trail. The audit trail was drafted based on the memos and the developed coding schemes. The authors evaluated the audit trail and agreed with research process.

## Findings

### What do teachers suggest as the most important content in genetics courses?

When we asked teachers to summarise what they hoped students would learn from upper-secondary school biology courses, the teachers mentioned different contents (Table 2, Table S1). We divided their answers to three distinct themes: 1) development of phenotype, 2) inheritance and continuity and 3) structure and function of the genes. Some teachers gave several descriptions that fitted two of these themes, but none described all three. The first theme, development of phenotype, contains descriptions that focused on understanding how genes and environment shape the development of different traits (i.e., genetic determinism). These descriptions were often related directly to how students themselves have developed and to their understanding of how human individuals have formed:

> Teacher J: “Humans are constructed by many factors, of which genome influences greatly, or they are things that we cannot influence ourselves; they come directly from the genome, but also genes do not dictate how we live our lives, what kind of persons we are, and how we behave.”

The second theme, inheritance and continuity, is centred on the concept that there is genetic continuity in the tree of life and that DNA copies itself from generation to generation. Teachers who described concepts relating to this theme saw the understanding of evolution as the focal point of whole field of biology and saw genetics as central to this understanding. Biodiversity was mentioned as one manifestation of this continuity. Sometimes the descriptions of the most important ideas were affective:

> Teacher C: “The common thread of life, from the beginning, the same genes are flowing; we are composed of genes from a million persons from thousands of years and then a new combination pops up, from the stream of life.”

The third theme, the function of the genes, was the simplest theme in terms teachers’ descriptions. They usually said that it was important to understand what genes are and how they function, while offering no reference or reason why it is so. Some teachers mentioned that in terms of general knowledge, it is important to know these topics.

> Teacher B: “If I say it concisely, what the gene is and how it functions is the core knowledge a student should have.”

### Use of human-related contexts in genetics teaching

#### GMOs

Most of the teachers approached ethical questions as being superimposed on the biological content within a course and they thought the students should know the biological contents of GMOs before discussing their ethical dimensions (Table S2). Some teachers also suggested that students have highly polarised opinions on GMOs before coming to a course and that “knowledge” could help them evaluate the different aspects of the debate.

> Teacher F: “We have two types of students, so that they are pretty black-and-white. Some of them have already been kind of brainwashed to think that “this is all great”, while a minority, or I don’t know if they don’t just dare to tell me, are against GMOs.”

Three teachers said that they use SSI to motivate students at the beginning of the course, while the other seven said that they first teach the biological content and then move to ethical discussions. Still, ethical questions were seen as secondary to biological content:

> Teacher J: “…there’s not always much time for discussions – the time spent in ethical discussions is always reduced from less than what is spent on the course texts.”

#### Human genetics

Teachers mentioned that the use of human examples in genetics is mostly limited to Mendelian disorders in the BI2 course (Table 2, Table S3) and more complex traits are then discussed at the end of BI2 or during the BI5 courses (Table 2, Table S4). Teachers commonly held the opinion that students are interested in hereditary phenomena in general (Table S5), but there is mismatch between how textbooks frame genetics and the students’ interests: while students are mainly interested in human genetics, the textbooks lack good examples and teachers did not feel themselves competent to go deeper into the topic:

> Teacher J: “… [a student asks] if I have blue eyes and my boyfriend has brown eyes, then what colour will our children’s eyes be, but unfortunately I have to try to contain their excitement as I don’t know the answers to their questions.”

Some teachers mentioned that they use classic, if not the most correct, examples like hair with a widow’s peak or the ability to roll the tongue. All teachers who used these examples said that, nevertheless, they mention to students that genetics is generally not that simple.

Two teachers mentioned that they try to avoid the human context in general and three other teachers said that they try to avoid discussing complex human traits, such as talent, intelligence, or human behaviour (Table 2, Table S6):

> Interviewer: Do you discuss how genetics affects learning? What if some students have genes that allow them to achieve better grades?
>
> Teacher E: No, no. (pause). No. Interviewer: No?
>
> Teacher E: No, we do not discuss that. Interview: No one is interested?
>
> Teacher E: No. I am not interested either (laughs)
>
> *Interviewer laughs*
>
> Teacher E: I think it is extremely sensitive issue. I would reconsider several times before discussing it.

Most teachers said that they discuss human behavioural genetics if students ask questions, but they do not introduce the topic themselves. In contrast, some teachers said the discussions are needed, especially in the context of racial issues:

> Teacher D: “It is relevant for the students if it is discussed in public, societal debate – [they may want to know if] citizens from certain continents are less intelligent than others–, and we have discussed these alternative news a lot, how they publish utter nonsense.”

### Controversial or sensitive issues in genetics

Half the teachers did not identify any sensitive or controversial issues thatthey would avoid (Table 2, Table S6). Among the other half there were differences in how they framed sensitive and controversial issues. Most of the argumentation was related to what is seen as biological general knowledge or avoiding misconceptions on genetics: teachers mentioned or implied that human traits are inherently so complex that there is a significant risk that students would form misconceptions on overtly genetic determinations of these traits. Furthermore, the lack of examples and lack of teacher competence was seen as leading to teaching without meaningful contents. The two teachers who avoid the topic humans as a context argued that humans are just one species, and it is not meaningful to concentrate too much on humans in biology.

Additionally, teachers mentioned that some issues cause discomfort for them or their students. Regarding students, some teachers acknowledged that discussing human heredity can pose several challenges (Table 2, Table S6). For example, using blood group testing can raise questions and even distress students if their blood group is not concordant with their parents’ blood groups. One teacher mentioned she does not want to do pedigrees on simple traits with students because of the “diversity of families” and she does not know the students’ backgrounds. The reason for describing these issues as “uncomfortable” was framed as a question of a teacher not knowing how to deal with discussing these issues or encountering unexpected reactions from the students. Those teachers willing to discuss genetic disorders of students or their families argued that generally those affected know best about the issues. One teacher also mentioned that sensitive issues bring up emotions, but that it is also natural in a classroom setting:

> Teacher C: “Sometimes I’ve gone and hugged a student – I find it a good way to calm down. – Every now and then I have tears in my eyes, but I think it’s important to show my own emotions in my teaching.”

### Teachers frame genetics teaching with different emphases

Teachers’ perceptions of the most important genetic content were closely related to their willingness to use a human context in their teaching or even what they said that students find interesting (Table 2). For example, none of the teachers whose theme in genetics teaching was classified as “development of phenotype” mentioned that they try to avoid human complex traits. In turn, both teachers who said that students are interested in gene testing had their theme grouped to “inheritance” and both teachers who mentioned students are interested in epigenetics to “development of phenotype.”

Three different general frameworks of teaching genes and their role arose from the analysis: *Developmental*, *Structural* and *Hereditary* (Table 3). We call these emphases of contents in genetics teaching, as they relate to how teachers argue not only for their choice of contents and which they see as the most important contents or themes, but also which contexts they use. They also reveal how they perceive student interest and whether they avoid certain topics. Furthermore, they align with perception of sensitive or controversial issues. We note that these emphases do not consider how teachers understand genes or their function, but rather what teachers see genetics to constitute from a teaching perspective.

**Table 3:** Three different content emphasis and the teacher perceptions and descriptions which differ between emphasis.

| **Content emphasis** | **Structural** | **Hereditary** | **Developmental** |
| --- | --- | --- | --- |
| *Central theme* | Gene structure and function | Continuity of DNA through time | Development of traits |
| *Human context* |  |  |  |
| 1. Human Mendelian disorders | Avoid | Use as examples | Use as examples |
| 2. Complex human traits | Avoid | If students ask | Use as examples |
| *Perceived student interest* | Monohybrid, dihybrid crosses | Gene tests, medical genetics | Epigenetics, complex human traits |

A *Developmental* emphasis frames the development of traits as the central theme in genetics and consequently teachers who used this approach were largely open to any discussions, they did not mention any topic they would avoid and most of them mentioned that they have regular discussions about complex human traits as they felt that students are most interested in these. Furthermore, teachers with *Developmental* emphasis were less likely to describe any perceived sensitive issues than other teachers (Table 2). These teachers were all comparatively the less experienced teachers of the interviewed group (12 or fewer years of experience). Their emphasis contrasts with two teachers who used a *Structural* emphasis mentioning only gene function as the central issue. These teachers mentioned avoiding discussing complex traits in humans or humans at all, as they found these both sensitive and not good examples of polygenic inheritance. In contrast, these teachers described hereditary analysis as an interesting part of the genetics course. They were among the most experienced teachers (> 20 years). A third emphasis, *Hereditary*, was characterised by emphasis on the continuity of DNA through the whole tree of life. This emphasis manifested in teachers’ answers as being somewhere between the two previous emphases. Teachers who used this emphasis were willing to discuss complex human traits if the students asked about them, but did not actively raise examples. They generally used an example of human skin colour as an example of a polygenic trait. More broadly, in genetics, they usually emphasised the understanding of phenomena related to DNA duplications, such as meiosis. A hereditary emphasis was used by both less and more experienced teachers.

In contrast to the issues involving the human complex traits, genetics content emphasis was not connected to how teachers taught GMOs (Table S2). While some teachers were more dismissive about teaching on SSI while discussing GMOs and said that there was not always time to go through those topics, they were not differentiated based on their genetics emphasis. Furthermore, one teacher who said that they use GMOs as a motivation at the beginning of the course to explain how genetics are important, said that they do not always have time to go into the ethics of GMOs. In general, lack of time is a general perception of teachers in different subjects and countries (Adams & Krockover, 1997; Archbald & Porter, 1994; Fuller, 1969) and the interviewed teachers also expressed this phenomena repeatedly. Most teachers mentioned lack of time and the need to closely follow the textbook were mentioned by most teachers, and noted that textbooks tend to discuss ethical dimensions of biotechnology at the end of the text (Aivelo & Uitto, 2015).

### Teacher self-identification and relationship to actual teaching

When we provided teachers with the descriptions of the genetics emphasis and our analysis of their interview, six teachers agreed with our analysis, three teachers disagreed and one teacher was unavailable (Table 2). All disagreeing teachers had their teaching emphasis labelled to Developmental. Two of the teachers also argued for that their emphasis was not the one they would have preferred for genetics teaching, but it was mostly dictated by national curriculum, which mentions evolution and development in different courses.

We obtained teaching diaries and other teaching materials with enough information for our analysis from five teachers. In categorization, the interrater reliability was high (Cohen’s kappa: 0.88). The concurrence between the interviews and the diaries was variable as some teachers were fully concordant (such as teachers B and H), whereas Teachers G and J had two discordant categorizations (Table S7). In total, 18 of 22 units were concordant between the interviews and diaries.

### Factors behind teachers’ choice of content

Although there were significant differences in teachers’ emphasis in genetics teaching, there were some similarities in their arguments on factors influencing their content choices (Table 2). While the teachers described their teaching in quite different terms, all except teacher E said that 1) they follow closely the textbook content. Because the contents of biology textbooks for upper-secondary schools is highly similar (Aivelo & Uitto, 2015), teachers clearly had different personal priorities on the most important contents and contexts. All schools follow 2) the national core curriculum (FNBE, 2004), and this was evident in teachers’ descriptions of their content choices (Table 2, 3). Furthermore, this was emphasised by the inclusion of GMOs in the biology core curriculum by all teachers (Table 3). The national core curriculum is also the basis for the tasks on the matriculation examination that the students take at the end of the upper-secondary school in Finland (Niemi et al., 2012) and teachers acknowledged that 3) previous exam questions guide their teaching.

The aforementioned factors were quite similar for each teacher, but our results also reveal perceived differences among teachers and the 4) school-specific circumstances in which they are working. Some of the teachers compared their school to other schools and suggested that some attributes of their school attract students with specific interests or motivation or competence to study biology. Likewise, teachers described differences in course arrangements, and noted whether it was possible to conduct experiments in the classroom. Furthermore, teachers expressed that there are 5) personal reasons that affect their course content selection. On many occasions the teachers acknowledged the limits of their competence, either regarding genetics contents, such as when they are unable to answer students’ complicated questions of the students, or pedagogically, when they mentioned they might have problems in successfully guiding classroom discussions.

## Discussion

### Teachers’ emphasis in genetics

Our findings suggest that there were fundamental differences in Finnish upper secondary school biology teachers’ perceptions on the most important themes in genetics and genetics teaching and subsequently how they chose course content and context while teaching genetics. The perceptions can be classified to three distinct areas of content emphasis, which we named *Structural*, *Hereditary* and *Developmental*. These emphases are formulated on the basis of how teachers interpret and use 1) the central themes in genetics, 2) human contexts in their genetics teaching, 3) students’ interest in different contents and contexts and 4) perceptions of sensitive or controversial issues in genetics.

Our findings are partly like those of Van Driel et al. (2007), who found separate subgroups of teachers who were either subject-matter oriented focusing on fundamental, theoretical concepts or learner-centred emphasising societal issues. While *Structural* emphasis can be seen as subject-matter oriented, *Developmental* emphasis is not learner-centred in similar sense as in the study by Van Driel et al., as the orientation is not as much societal as it is personal. Stewart, Cartier and Passmore (2005) outlined three different models of genetics understanding: inheritance pattern, meiotic and biomolecular. These models are quite close to our concept of content emphasis: inheritance patterns model and *Hereditary* emphasis are similar. *Structural* resembles the meiotic model while *Developmental* has less in common with the biomolecular model.

The diversity of emphases can be explained by the complex educational context in the Finnish upper secondary school, which aims not only to train students for tertiary education, but also to develop scientific literacy in those students who may not study biology further. The biology course “Cells and heredity” is compulsory for all students of which approximately one-third complete the biology part of the matriculation examination (Matriculation Examination Board, 2019). Thus, the teachers balance what Roberts (2007) referred to as vision I (as knowledge within science) and vision II (as knowledge in everyday situations) of scientific literacy.

Moreover, genetics content emphasis can be seen as partly overlapping with “science teaching orientations” (Magnusson, Krajcik, & Borko, 1999) as they contain knowledge of the importance of different concepts, interpretation of curricula, the motivations of the students, and representations and context of core concepts. In comparison, while the science teaching orientations describe teachers’ perceptions about teaching and especially instruction methods, we did not find that genetics content emphasis would limit the instruction methods. Nonetheless, the different emphases rises the question of how differential teacher understanding of core concepts and contexts influences teaching methods or orientations to teaching science. In a follow-up study, we aim to study teachers’ gene concepts and whether they relate to a different emphasis.

### Human-related contexts involve controversial and sensitive issues

Our research setting in comparing three different human-related contexts - GMOs, Mendelian human traits and complex human traits - contrasts the effects of curriculum-dictated context choice and free choice by teachers, and highlights the difference between personal and societal relevance. Our interviews showed a paradoxical approach by teachers: while they said that genetics is a societally relevant topic, and that students should learn analytical tools to take part in decision-making and be responsible consumers, this was not evident in their descriptions of their teaching. Without exception, teachers formulated the basic science as the main issue and, in many cases, societal aspects of GMOs were described to be taught only “if there was time at the end of the course.” Our results agree with Tidemand and Nielsen’s (2017) suggestion that emphasis on biological content (as opposed to more societal context) is driven by teachers’ identity as biology teachers (see also Pedretti, Bencze, Hewitt, Romkey, & Jivraj, 2008). Nevertheless, all teachers did teach GMOs as they are explicitly mentioned in national curriculum.

In comparison, some teachers described avoiding human genetics contexts that could be seen as personally highly relevant to students. These teachers were also more likely to describe controversial or sensitive issues related to genetics teaching. It is noteworthy, that teachers framed sensitive issues in human genetics in relation to students personally, as something that concerns individual students and not so much the society at large. Thus, in Rowling’s (1996) categorization, the teachers were more worried about sensitive issues than controversial issues. Furthermore, this suggests that there is a trade-off between personal relevance to students and perceived problems arising from sensitive issues. As this was not mentioned in relation to GMO teaching, teachers seem to be more hesitant about this personal relevance, rather than societally controversial issues.

The avoidance of human contexts is a complex issue as teachers used numerous reason for steering clear of topics in human genetics context, for example: a) students are not interested in these topics, b) teachers do not have enough content knowledge, c) teachers do not have pedagogical knowledge for teaching sensitive issues and d) courses discussing genetics in human context would lead to negative learning outcomes, such as misconceptions. In general, controversial issues were thought to lead to misconceptions, whereas sensitive issues were seen to lead to awkward situations for individual students. We are not able to assess how relevant these various factors are, but it is clear from our content emphasis classification that there are fundamental differences in how teachers perceive the most important contents and contexts in genetics. Furthermore, contrary to the previous studies (Hess, 2008; Phillips, 1997), our study suggests that Finnish teachers are open to discuss complex human traits and other sensitive issues in classroom even when they are not experienced.

### Limitations of the study

While the number of interviews in our study is limited, we reached data saturation rapidly. One reason for this may be the similarities in the educational background of the teachers, as all of them had a master’s degree with biology as the major subject, and pedagogical studies in teacher education as a minor subject. Moreover, the textbooks used by the teachers are quite similar, emphasizing gene structure and function (Aivelo & Uitto, 2015). Due to the small number of participants and limited knowledge on teachers’ background, the generalizability of this study is limited and we cannot discuss other factors than those mentioned by the teachers: namely, the role of the chosen textbooks, biology curriculum, practicing for the matriculation examination and school-specific and personal factors. The interpretive disagreements on genetics content emphasis between some teachers and researchers were all related to a *Developmental* emphasis. Nevertheless, the interrater reliability in categorization and the concordance between teacher interviews and diaries was rather high.

Potential sources of bias were minimized by allowing the interviews to be as freely advancing as possible and questions were designed to prevent confirmation bias by probing for disconfirming answers and leading questioning by beginning each strain of questions with general questions. Research positionality was reflected regularly through interactions between the authors and in discussions with outside researchers and biology teachers. The authors have multi-faceted relationship towards participant community as they are involved in teacher education and in-service teachers’ continuing education and both have backgrounds as upper-secondary school biology teachers. Both authors have also been involved in the national core curriculum process. Thus, the authors are insiders in the participant community but also hold positions of power. This setting was approached by emphasizing to teachers that they are experts in teaching practice and that the researchers were genuinely interested in their answers.

### Implications for research and teaching practice

The national curriculum for upper-secondary schools gives substantial freedom to teachers to interpret the contents and goals of biology education in classroom practice (FNBE, 2004; Niemi et al., 2012). This may partly explain the fundamental differences in content emphasis that we found. Consequently, in school practice, teacher education and in-service training, the teachers should be made more aware and provide opportunities for self-reflection on the emphasis they take in teaching science.

We also suggest that our findings on which contents teachers choose for their teaching provides a well-grounded hypothesis for further research on the content perspective of experienced, autonomous biology teachers. The relationship between content emphasis and the choice of course content could provide a more widely applicable hypothesis for studying teaching and learning genetics in biology education, because teachers have freedom to choose whether to apply a sensitive human context to their teaching.

Considering GMO SSIs being integrated in the teaching, independent of teacher inclinations, we suggest that curriculum development would be a valuable tool if genetics education aims to better incorporate societal and personal relevance. Furthermore, curriculum development needs to relate to teacher education emphasizing pedagogical content knowledge (Käpylä, Heikkinen, & Asunta, 2009). While teachers appreciate a “knowledge first” approach to SSI and avoidance of human related topics, there is a perceived lack of useful and tested teaching materials. Content knowledge is important for successful reasoning regarding SSI: thus, a delicate balance needs to be sought (Lederman et al., 2014; Sadler & Zeidler, 2004)

From our interviews, it is clear that personal relevance in teaching can have various effects in the classroom: while some teachers see it as a possibility, some seem to avoid it for a number of reasons. This topic needs to be addressed more in professional development. In general, the ways of teaching controversial and sensitive issues are not well-studied and the recommendations themselves are controversial (Christopher Oulton et al., 2004). Thus, both societal and personal relevance should be taken more into account in science and especially in biology teacher education and in-service training.

## Conclusions

Based on qualitative case study and teacher interviews, we have found that teachers’ perceptions of genetics teaching reflected three different emphases that we named as *Structural*, *Hereditary*, or *Developmental* content emphasis. These emphases consist of teachers’ perceptions of the most important themes in genetic content, willingness to teach about human traits, perceived sensitive or controversial issues in genetics and students’ perceived interests. Interestingly, teachers having *Structural* emphasis described avoidance of a human genetics context in their teaching, while teachers with a *Developmental* emphasis described very abundant use of human genetics contexts. Thus, teachers’ perceptions of which themes in genetics are the most important could also shape how likely they are to use personally relevant contexts in their teaching. As we did not observe the actual teaching practice, we do not know how well these emphases manifest in teaching itself and whether these have an actual effect on student learning outcomes. Our ongoing research project could shed light on this by comparing student interests and attitudes to their teacher’s genetics content emphasis. Nevertheless, we suggest that teachers’ perceptions on the most important themes in their teaching can have wide-ranging consequences, for example, inclusion of socioscientific issues in the teaching.

Our results also revealed different approaches to the sensitive and controversial issues in genetics teaching. Not all teachers perceived that sensitive or controversial issues would affect their teaching and those who did, usually described sensitive rather than controversial issues, thus suggesting that teachers are more worried about personal issues in genetics. Indeed, sensitivity was sometimes used as a justification to exclude contents or contexts that are personally relevant to genetics teaching. We note that teachers would need more support to handle controversial and sensitive issues in the classroom. In contrast to personally relevant human genetics, the Finnish curriculum specifically mentions GMOs and compels teachers to discuss them. Subsequently, every teacher mentioned that they discuss GMOs. Thus, we also argue that curricular development is an effective way to increase the prominence of societal or personal relevance in biology education.

## Acknowledgments

The authors wish to thank the ten biology teachers who participated in this study, Marlene Broemer for checking the language in manuscript, Lotta Aarikka for transcribing the interviews and providing debriefing during initial analysis, Justus Mutanen for teaching diary analysis and fruitful discussions and Niklas Gericke for comments on an earlier version of the manuscript.

## Appendix

Supplemental material in Figshare includes following tables containing representative quotes from teacher interviews:

Table S1: Teacher’s descriptions of the central theme of their teaching.

Table S2: Teacher’s descriptions of GMOs in their teaching.

Table S2: Teacher’s descriptions of the human Mendelian disorders in their teaching.

Table S3: Teacher’s descriptions of the complex human traits in their teaching.

Table S5: Teacher’s descriptions of the perceived student interest in genetics.

Table S6: Teacher’s descriptions of perceived sensitive issues.

Table S7: The concurrence between teaching diary and other materials in comparison to teacher interviews.

